# The predicted acetoin dehydrogenase pathway represses sporulation of *Clostridioides difficile*

**DOI:** 10.1101/2023.07.28.551048

**Authors:** Daniela Wetzel, Arshad Rizvi, Adrianne N. Edwards, Shonna M. McBride

**Affiliations:** Department of Microbiology and Immunology, Emory University School of Medicine; Emory Antibiotic Resistance Center, Atlanta, GA, USA

**Keywords:** *Clostridioides difficile*, *Clostridium*, acetoin, pyruvate, pyruvate dehydrogenase, *aco*, metabolism, sporulation

## Abstract

*Clostridioides difficile* is a major gastrointestinal pathogen that is transmitted as a dormant spore. As an intestinal pathogen, *C. difficile* must contend with variable environmental conditions, including fluctuations in pH and nutrient availability. Nutrition and pH both influence growth and spore formation, but how pH and nutrition jointly influence sporulation are not known. In this study, we investigated the dual impact of pH and pH-dependent metabolism on *C. difficile* sporulation. Specifically, we examined the impacts of pH and the metabolite acetoin on *C. difficile* growth and sporulation. We found that expression of the predicted acetoin dehydrogenase operon, *acoRABCL*, was pH-dependent and regulated by acetoin. Regulation of the *C. difficile aco* locus is distinct from other characterized systems and appears to involve a co-transcribed DeoR-family regulator rather than the sigma^54^-dependent activator. In addition, an *acoA* null mutant produced significantly more spores and initiated sporulation earlier than the parent strain. However, unlike other Firmicutes, growth and culture density of *C. difficile* was not increased by acetoin availability or disruption of the *aco* pathway. Together, these results indicate that acetoin, pH, and the *aco* pathway play important roles in nutritional repression of sporulation in *C. difficile*, but acetoin metabolism does not support cell growth as a stationary phase energy source.

**IMPORTANCE:** *Clostridioides difficile,* or *C. diff*, is an anaerobic bacterium that lives within the gut of many mammals and causes infectious diarrhea. *C. difficile* is able to survive outside of the gut and transmit to new hosts by forming dormant spores. It is known that the pH of the intestine and the nutrients available both affect the growth and sporulation of *C. diffiicile,* but the specific conditions that result in sporulation in the host are not clear. In this study, we investigated how pH and the metabolite acetoin affect the ability of *C. difficile* to grow, proliferate, and form spores. We found that a mutant lacking the predicted acetoin metabolism pathway form more spores, but their growth is not impacted. These results show that *C. difficile* uses acetoin differently than many other species and that acetoin has an important role as an environmental metabolite that influences spore formation.

## INTRODUCTION

*Clostridioides difficile* is an anaerobic, toxin-producing gastrointestinal pathogen that is the leading cause of antibiotic associated diarrhea (1). As a strict anaerobe, the ability of *C. difficile* to form spores is critical for both its survival outside of the host and the transmission of the pathogen to new hosts (2). While the formation of *C. difficile* spores appears morphologically similar to other endosporulating bacteria, the environmental cues and mechanisms that lead to sporulation of *C. difficile* differ from other spore-forming species and are not well defined (3–6).

Two conditional factors that greatly influence *C. difficile* spore formation are nutrient availability and the pH of the surrounding environment (3, 7). Prior work demonstrated an acute effect of pH on the sporulation of *C. difficile*; however, the mechanism by which pH affects sporulation is not understood (7). We hypothesized that pH influences the initiation of sporulation in *C. difficile* by changing the availability and metabolism of important nutrients. One important nutrient that is metabolized in response to pH in many bacteria is the intermediate metabolite, acetoin (3-hydroxy-2- butanone). During active growth, bacteria decrease the environmental pH due to the production and export of acidic products of glycolysis (8). This drop in pH stimulates the synthesis of acetoin from pyruvate, which is then secreted from the cell, reducing the acidification of the medium (9).

As preferred nutrients are depleted during stationary phase growth, acetoin is imported and catabolized (8). Acetoin metabolism is performed by Acetoin Dehydrogenases (AoDH), which are enzyme complexes comprised of subunits E1α (AcoA), E1β (AcoB), E2 (AcoC), and E3 (AcoL) (10–12). In the model sporulating bacterium, *Bacillus subtilis*, the metabolism of acetoin during post-exponential growth supports spore formation (13, 14). Although acetoin synthesis and degradation has been examined in clostridial species as it relates to industrial fermentation, food science, and solvent production, the relationship between pH, acetoin metabolism, and sporulation is not clear (10, 15–18) In this study, we sought to determine if acetoin availability and utilization contributes to *C. difficile* spore formation. Herein, we investigated the predicted acetoin metabolic gene cluster (*acoRABCL, CD0035-CD0039*) and assessed the impact of this pathway on *C. difficile* gene expression, growth, and sporulation. We observed that expression of the *aco* operon was pH-dependent and subject to feedback regulation by acetoin. Further, we found that a mutant in the predicted acetoin metabolic pathway produced significantly more spores than the wild-type strain, but surprisingly, acetoin metabolism had no apparent effect on growth. These observations suggest that acetoin plays an important role in nutritional repression of sporulation in *C. difficile*, but does not function as an energy storage vehicle that can support post-exponential growth and cell proliferation.

## RESULTS

### Expression of the *aco* locus is pH-dependent

In previous work, we observed that sporulation of *C. difficile* substantially increased as the pH of the growth medium was raised, which corresponded with considerable changes in gene expression. In an effort to determine the mechanisms involved in pH-dependent sporulation, we identified transcripts that were differentially expressed under low and high relative pH conditions. Of these, the predicted acetoin dehydrogenase genes demonstrated significantly decreased expression that strongly correlated with increases in the pH of the medium (**Figure 1; Figure S1**). During growth on sporulation agar, we observed a ∼5-fold decrease in *acoA* transcription as the pH increased from 5.5 to 6.5, ∼8-fold decrease at pH 7.5 (relative to pH 5.5), and a greater than 10-fold decrease in transcription at pH 8.5. Hence, *aco* expression is greatest at low pH and decreases with corresponding increases in the pH of the medium. Based on these data and prior work showing an effect of acetoin on sporulation in *B. subtilis* (14), we hypothesized that utilization of acetoin in *C. difficile* may contribute to the observed pH-dependent sporulation phenotypes.

**Figure 1.**
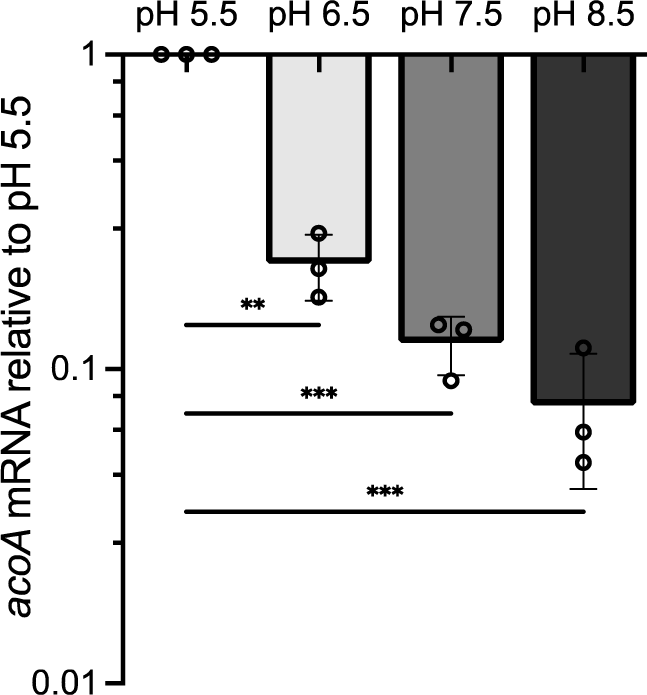
Expression of the *aco* locus is pH-dependent. qRT-PCR analysis of *CD0036* (*acoA*) expression in *C. difficile* strain 630Δ*erm* grown 12 h on 70:30 agar at pH 5.5, 6.5, 7.5, and 8.5, normalized to pH 5.5. The means and standard deviation of the mean for three independent replicates are shown. Expression levels were analyzed by one-way ANOVA and Dunnett’s multiple comparisons test compared to pH 5.5. Asterisks indicate *P* values: * ≤ 0.05; ** ≤ 0.01; *** ≤ 0.001

### Defining the acetoin metabolism locus

To identify effects of acetoin metabolism on *C. difficile,* we first examined the predicted acetoin utilization gene cluster for its transcriptional composition. The *aco* gene cluster was annotated to include *acoA, acoB, acoC,* and *acoL* (**Figure S1**; *CD0036-CD0039*). The tandem arrangement of *acoABCL* in the *C. difficile* genome is consistent with the organization of the acetoin locus of *B. subtilis.* However, an additional ORF, *CD0035,* is present upstream of *acoA* and is predicted to encode a DeoR-family transcriptional regulator, whereas *B. subtilis* encodes a sigma 54-dependent accessory activator that is transcriptionally separate and downstream of *acoABCL* (**Figure S1**). To define the transcriptional units within this cluster, we examined the entire *aco* locus for transcript linkage. Using cDNA generated from logarithmic cultures, we performed PCR to amplify across adjacent open reading frames (**Figure S2**). Products were amplified across all ORFs from *acoR* to *acoL* (*CD0039*), but not between *acoL* and the downstream ORF, *CD0040*. Accordingly, we designated *CD0035* as *acoR* to reflect the inclusion of the regulator within the *aco* operon.

### Disruption of *acoA* results in greater sporulation of *C. difficile*

In the model spore-former *B. subtilis,* acetoin reduces spore formation and sporulation is further decreased when the acetoin pathway is disrupted (14). To determine if acetoin utilization impacts spore formation or pH-dependent phenotypes in *C. difficile*, we generated a deletion mutant in the predicted *acoA* gene, which is required for acetoin pathway function (14) (**Figure S3**). The resulting mutant (MC1719) was examined for the ability to produce ethanol-resistant spores on sporulation agar under a range of pH conditions. Relative to the parent strain, the *acoA* mutant demonstrated considerably greater spore formation, which was statistically significant at both pH 6.5 (6.9-fold) and pH 7.5 (2.8-fold) (**Figure 2**). The hypersporulation of the *acoA* mutant was fully rescued by chromosomal complementation with the *acoRABCL* operon (**Figure S4)**. Further, the addition of acetoin to the medium did not consistently alter sporulation in the wild-type or *acoA* mutant strain. These data suggest that the metabolism of acetoin either reduces the ability of the bacteria to engage the sporulation pathway or reduces mature spore production.

**Figure 2.**
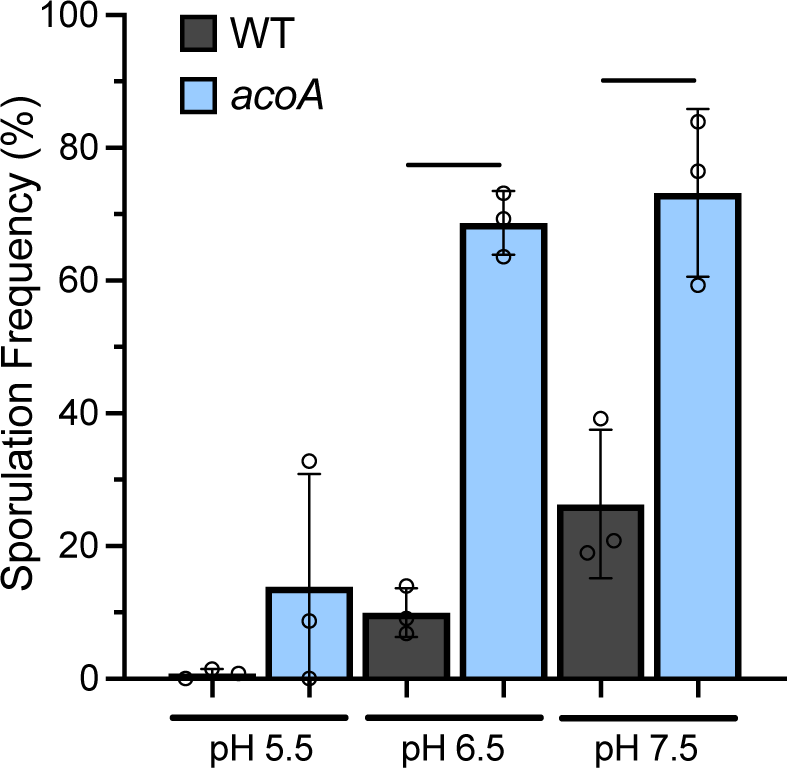
The Aco operon reduces spore formation across a range of pH conditions. Ethanol-resistant spore formation of wild-type (630Δ*erm*) and the *acoA* mutant (MC1719) grown on 70:30 sporulation agar at pH 5.5, 6.5, or 7.5 for 24 h. Data represent the percentage of spores relative to total viable cells at 24 h, shown with the standard deviation for three independent replicates. A Student’s two-tailed *t* test was used to compare the 630Δ*erm* and *acoA* mutant outcomes for each pH condition. **** indicates *P* value ≤ 0.0001; ** indicates *P* value ≤ 0.01

We further investigated the impact of the *aco* locus on the initiation of sporulation. To do so, we fused the promoter of the early sporulation locus, *spoIIG*, to the alkaline phosphatase (AP) reporter, *phoZ*, and examined activity over time during growth in sporulation broth cultures. The *spoIIG* operon encodes the early mother cell sigma factor, SigE, which is transcribed upon phospho-activation of the master sporulation regulator, Spo0A (19–21). During logarithmic phase growth, the *acoA* mutant and parent strains displayed similarly low levels of expression from the *spoIIG* promoter, signifying negligible initiation of sporulation within the population (**Figure 3**). But, by the onset of stationary phase (T_0_), the *acoA* mutant demonstrated 9-fold greater reporter activity than the parent strain, indicative of robust sporulation-specific gene expression that precedes the initiation timing observed for the wild-type strain. These results suggest that it is the stationary phase utilization of acetoin which suppresses initiation of the sporulation pathway, rather than the production or presence of acetoin.

**Figure 3.**
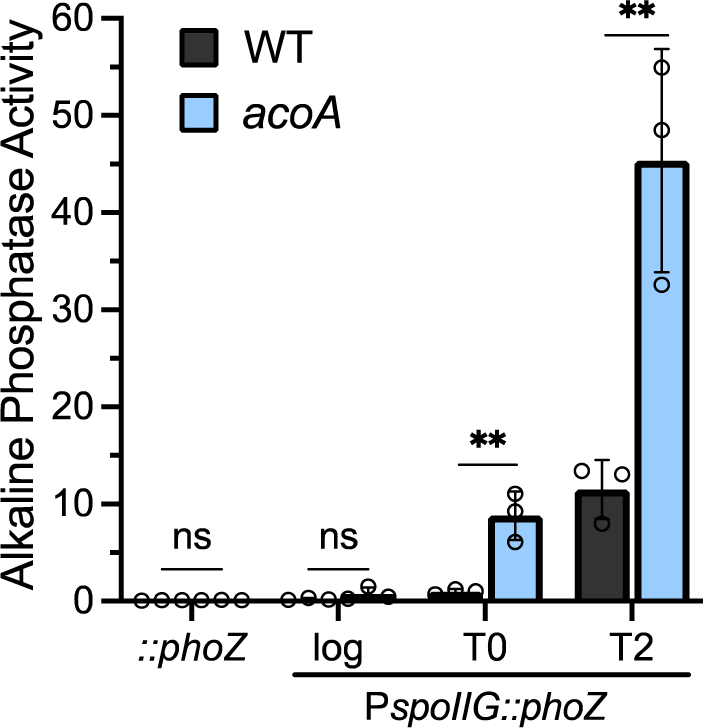
The Aco operon delays sporulation. Alkaline phosphatase (AP) activity of the early sporulation P*spoIIG*::*phoZ* reporter fusion in *C. difficile* strain 630Δ*erm* (MC492) and the *acoA* mutant (MC1760), grown in 70:30 sporulation medium containing 1 µg/ml thiamphenicol. Samples were assayed from logarithmic growth (OD_600_ 0.5), early stationary phase (OD_600_ 1.0; T0), and two hours after the onset of stationary phase (T2). The promoterless *phoZ* reporter carried by the parent strain (MC448, 630Δ*erm*) and *acoA* mutant (MC1759) were included as negative controls. The means and standard deviations for three independent replicates are shown. An unpaired Student’s *t*-test was used to compare the activity in the *acoA* mutant to the parental strain, per timepoint. ** indicates *P* value ≤ 0.01

### Acetoin metabolism has limited effects on growth or toxin production in *C. difficile*

A key trigger for sporulation is limitation of nutrients in the environment (22–25). Similarly, the nutrient limiting conditions that favor the sporulation of *C. difficile* generally support toxin production. The concurrent production of toxin and spores occurs due to an overlap of regulatory factors that control both processes, including CcpA, CodY, RstA, and Spo0E (3, 25–30). Considering the increased sporulation of the *acoA* mutant, we examined the impact of the locus on toxin production. To this end, wild-type and the *acoA* mutant strain were cultivated in TY medium and toxin TcdA and TcdB levels in the supernatant were assessed by ELISA after 24 h of growth (**Figure S5**). While there was a trend for the *aco* mutant to generate more toxin, the differences did not reach statistical significance (*P*= 0.051).

The production and degradation of acetoin can independently impact the environmental pH, cellular redox balance, and growth of bacteria (11). To determine if *C. difficile* uses acetoin as a carbon source to support cell growth and replication, we assessed growth of the wild-type and mutant strains in a minimal medium with and without acetoin (**Figure 4**). No significant differences in growth or pH were observed between the *acoA* mutant and parent strain, with or without the addition of acetoin. However, the addition of 30 mM acetoin did decrease the overall growth and culture density by ∼20% for both strains (**Figure 4AB**), suggesting that excess acetoin has deleterious effects on cell proliferation. Similar, but respectively lesser, effects were observed with the addition of acetoin to the medium at 10 mM and 20 mM (not shown). These growth results are in contrast to those seen for *B. subtilis*, which demonstrates increased proliferation with the addition of acetoin to wild-type cultures and can use acetoin as a sole carbon source (14, 31). Moreover, disruption of the acetoin pathway in *B. subtilis* resulted in reduced growth for an *acoA* mutant, even in the absence of exogenous acetoin (31). The lack of demonstrable growth impact of acetoin catabolism in *C. difficile* suggests that acetoin may play a lesser role as a carbon storage molecule in this bacterium than in other Firmicutes (Bacillota).

**Figure 4.**
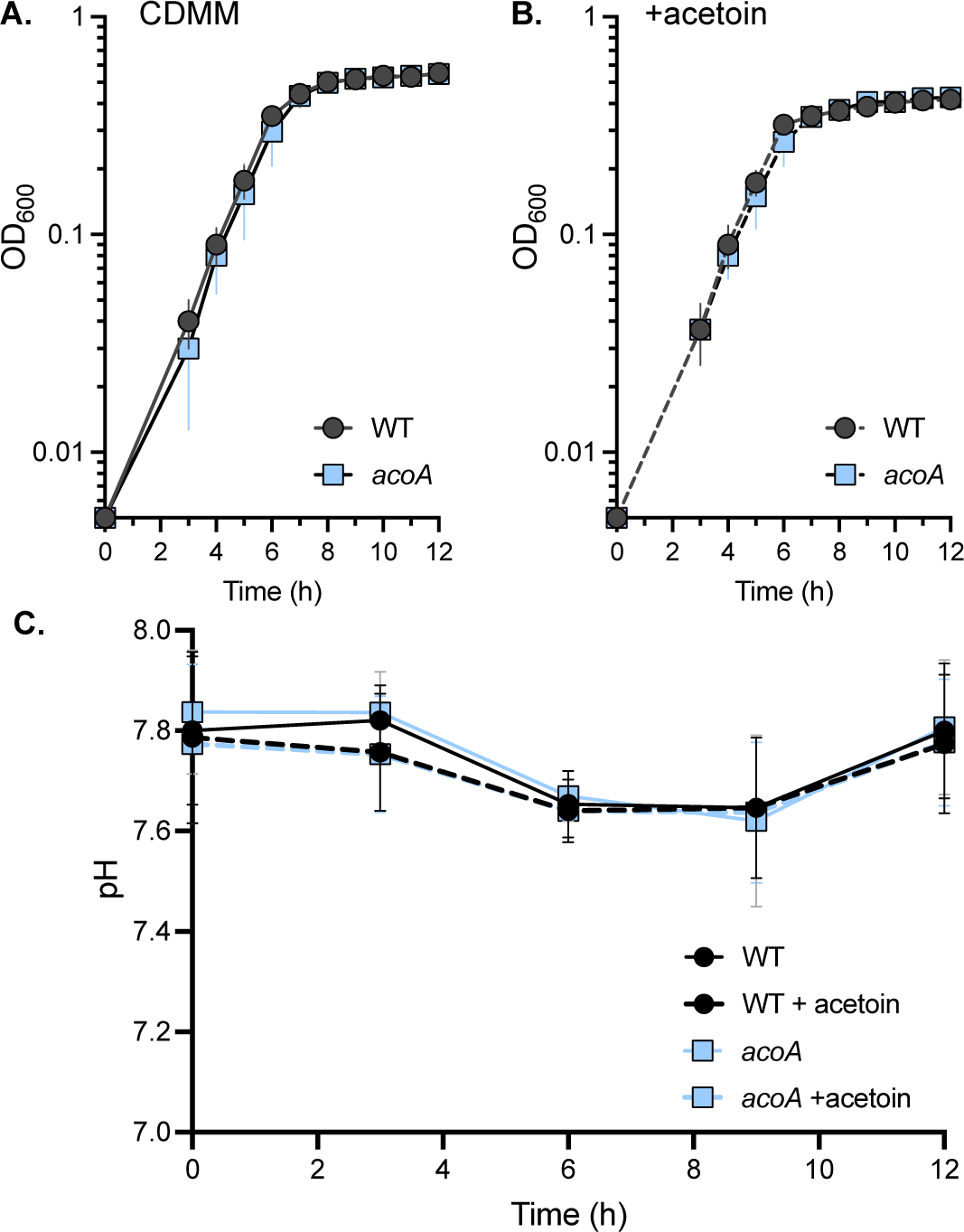
Growth and pH are not impacted by the disruption of acetoin metabolism. Strain 630Δ*erm* (WT) and the *acoA* mutant (MC1719) were cultivated in **A)** CDMM or **B)** CDMM with 30 mM acetoin. **C)** pH measured from the same cultures. Graphs are plotted as the means +/- SD from three independent replicates. Differences in the means values of WT and *acoA* at each time point and between the same strain +/- acetoin were analyzed by two-way ANOVA with Tukey’s post-hoc test; no significant differences in growth or pH were observed between the strains in either condition. Decreased growth with acetoin were noted for both strains between H7-H12 (*P* ≤ 0.05).

### Expression of the *aco* operon is repressed by pyruvate and acetoin feedback

Despite the similarity of the *C. difficile* AcoABCL amino acid sequences to orthologs of other Firmicutes, the predicted AcoR regulator of *C. difficile* bears no significant resemblance to other characterized AcoR regulators (**Figure S1**). The previously characterized AcoR regulators are sigma 54-dependent accessory activators that are often transcribed independently of the acetoin metabolism genes (11, 32–34). Unlike other *aco* regulatory factors, the *C. difficile* AcoR is a DeoR-family transcriptional regulator encoded within the same operon as the *aco* metabolic genes. In addition, the *aco* locus of *C. difficile* does not appear to have sigma^54^/SigL regulatory elements or dependence (35, 36). The *aco* genes of other Gram-positive bacteria are inducible by pH and acetoin, and are also subject to regulation by carbon catabolite repression through the global regulator, CcpA (14, 33, 37). Though the *C. difficile aco* locus is regulated by pH (**Figure 1**), no CcpA or glucose-dependent regulation has been observed for the *C. difficile aco* operon, nor is a characteristic *cre* site present (24). Thus, regulation of *C. difficile aco* gene expression does not appear to be under the same regulatory influences as previously characterized systems.

To understand how this pathway and the associated sporulation phenotypes are controlled, we examined expression from the predicted *aco* promoter under different conditions. For this, we generated a transcriptional fusion of the predicted promoter region upstream of the *acoRABCL* operon to the *phoZ* reporter (P*aco::phoZ*) and expressed the resulting plasmid in the wild-type and *acoA* strains. In contrast to other species, expression from the *aco* promoter was repressed by acetoin in wild-type *C. difficile*, rather than induced (**Figure 5**). P*aco::phoZ* activity was reduced in the *aco* mutant relative to the parent strain, which further supports the contribution of acetoin to feedback regulation. The addition of pyruvate to the cultures also resulted in significant reduction in promoter activity, indicating transcriptional repression by this precursor metabolite. However, the mechanism for this regulation is not clear.

**Figure 5.**
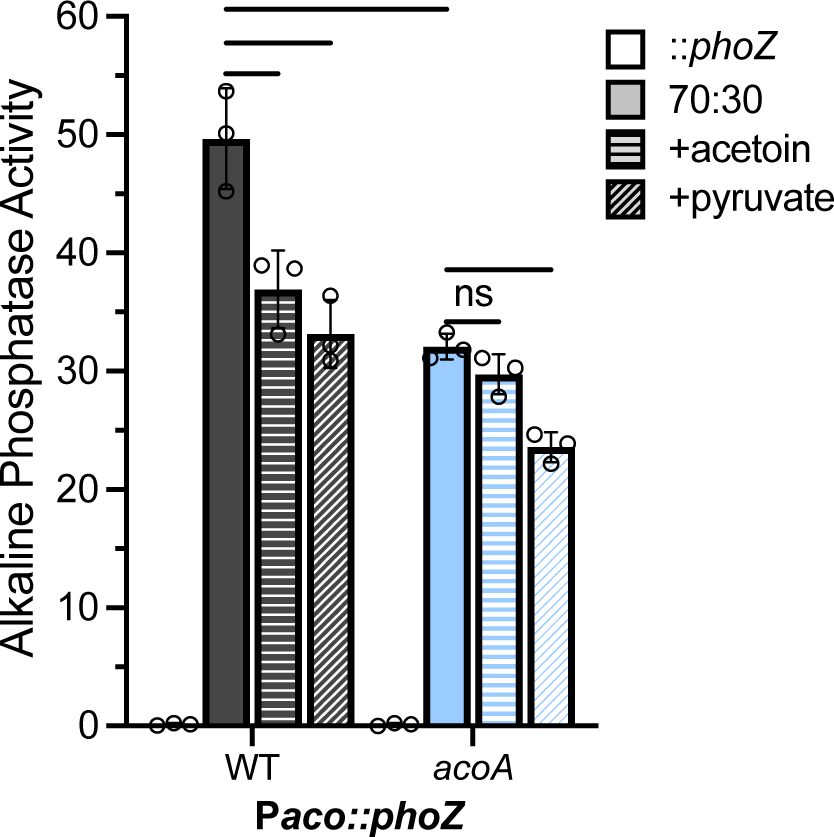
Expression of the acetoin operon is impacted by metabolites and function of the acetoin pathway. Alkaline phosphatase (AP) activity of the P*aco*::*phoZ* reporter fusions in *C. difficile* strain 630Δ*erm* (MC1827) and the *acoA* mutant (MC1828), grown in 70:30 medium containing 1 µg/ml thiamphenicol, with or without 30 mM acetoin or pyruvate. Samples were taken during logarithmic growth (OD_600_ 0.5). The promoterless *phoZ* reporter carried by the parent strain (MC448, 630Δ*erm*) and *acoA* mutant (MC1759) were included as negative controls. The means and standard deviations for three independent replicates are shown. A two-way ANOVA with Tukey’s post-hoc test was used to compare the activity of strains by growth condition. Asterisks indicate *P* values: * ≤ 0.05; ** ≤ 0.01; *** ≤ 0.001; **** ≤ 0.0001.

Acetoin dehydrogenase systems (AoDHs) are large, lipoate-dependent enzyme complexes that share high sequence similarity to pyruvate dehydrogenases (PDH) [19– 21]. To verify the functions of the enzymes encoded by the *acoABCL* genes, we performed modified Voges-Proskauer tests to assess the utilization and production of acetoin by the wild-type and *acoA* mutant (38). As shown in **Figure S6**, no significant difference in acetoin concentration was observed between wild-type *C. difficile* (strain 630Δ*erm*) and the *acoA* mutant grown in BHIS medium with 0.5 mM acetoin. Additional tests were performed to assess acetoin production at multiple growth stages and with the addition of precursor substrates glucose and pyruvate; however, no acetoin was detectable by our assays under the conditions tested (not shown). These results cannot rule out acetoin production or metabolism under other growth conditions, including within the host.

## DISCUSSION

Spore formation is a critical process for *C. difficile* and other anaerobes that allows the bacteria to survive in atmospheric oxygen and spread in the environment. Although sporulation is essential, it is also energy intensive and represents a state of dormancy that requires specific conditions to escape. As a result, all spore-forming bacteria have evolved regulatory mechanisms to prevent the initiation of sporulation when conditions can support growth and replication. The availability of nutrients and the pH of the environment are major factors that limit the entry into sporulation (3, 7). In this study, we examined the impact of the metabolite acetoin to determine if this nutrient is important to post-exponential growth and sporulation in *C. difficile,* as it is for other organisms. We found that while predicted acetoin metabolism locus plays a significant role in the production of *C. difficile* spores and the initiation of sporulation, there are considerable differences in how this bacterium reacts to acetoin, the phenotypes associated with acetoin metabolism, and the regulation of this pathway, compared with other species.

In most bacteria, including the model spore-former *B. subtilis,* acetoin acts as an important energy storage molecule that supports post-exponential growth and cell proliferation (14). Surprisingly, the addition of acetoin to the medium did not change the apparent growth rate or final cell density of *C. difficile* cultures and added acetoin had no significant effects on sporulation (**Figure 3, S4**). However, loss of *acoA* increased spore formation. Also, despite the pH-dependent expression of the *aco* operon, disruption of *acoA* had no significant impact on the pH of the medium (**Figure 1, 4**). These findings suggest that acetoin is not a preferred carbon source for growth, but is sufficient to delay sporulation initiation (**Figure 3**). Considering that acetoin can be reduced or oxidized to 2, 3-butanediol, it is possible that acetoin utilization serves more as a mechanism for balance of redox for *C. difficile* than as a significant source of energy or a mechanism for reducing acidification of the environment(38). Redox balancing contributes to both toxin production and sporulation; however, the specific connections between these processes has not been determined (39, 40).

In addition to differences in acetoin growth and sporulation phenotypes, the regulation of the acetoin metabolic genes in *C. difficile* is unlike that of previously characterized systems (10, 11, 31, 33, 34). In contrast to other bacteria, expression of the *aco* operon is not induced by acetoin in *C. difficile*, but rather *aco* gene expression is reduced in the presence of acetoin and the precursor metabolite, pyruvate (**Figure 5**). The most obvious cause of these differences in regulation is that the AcoR regulator of *C. difficile* is a co-transcribed DeoR-family regulator, rather than the sigma^54^-dependent activator that regulates transcription of the *aco* genes in other species (**Figure S1)** (11, 33, 41). DeoR-family regulators often bind directly to sugars, which act as effectors and control regulator function, while sigma^54^-dependent genes respond to nitrogen availability (42). It is likely that the *C. difficile* AcoR binds to a metabolic effector(s), which would result in more direct regulation, rather than regulation by general nutrient depletion that occurs with other AcoR regulators. Further, acetoin metabolism of many characterized species are subject to carbon catabolite repression (CCR), which restricts the utilization of lower quality metabolites when optimal nutrients are available (14, 43). Based on published information, the regulation of *aco* transcription in *C. difficile* does not appear to be dependent on the characterized metabolic-associated regulators CodY, CcpA, SigH, SigB, SigD, SigL, Spo0A, Rex, PrdR, or ClnR (19, 24, 26, 27, 35, 36, 39, 44–47). However, further studies are needed to characterize AcoR involvement and potential mechanisms of *aco* transcriptional regulation.

These results demonstrate both the conditional expression of the predicted acetoin metabolic genes and that the *aco* operon has a significant impact on *C. difficile* spore formation. The contrasts in acetoin utilization and regulation suggest that the differences in niches and nutritional requirements of *C. difficile* have influenced the utility of acetoin as an energy source. Further study is needed to determine the mechanism of AcoR regulation in *C. difficile* and the role of acetoin metabolism in pathogenesis.

## MATERIALS AND METHODS

### Bacterial strains and growth conditions

Bacterial strains and plasmids used in this study are listed in **Table 1**. An anaerobic chamber (Coy Laboratory Products) was used to cultivate *C. difficile* at an atmosphere of 10% H_2_, 5% CO_2_, and 85% N_2_ at 37°C, as previously described (48, 49). Strains were routinely grown fresh from -70°C stocks on brain heart infusion agar supplemented with yeast extract (50) (BHIS; Βecton Dickinson Company) broth or agar plates or TY medium (48) in the presence of 1-5 µg/mL thiamphenicol (Sigma-Aldrich) or anhydrotetracycline (ATc) when needed. *Escherichia coli* was grown at 37°C in LB medium (51) with 100 μg/mL ampicillin and/or 20 μg/mL chloramphenicol (Sigma-Aldrich) as indicated. Following conjugation with *C. difficile*, *E. coli* was counterselected using 100 µg/mL kanamycin (52).

**Table 1.**
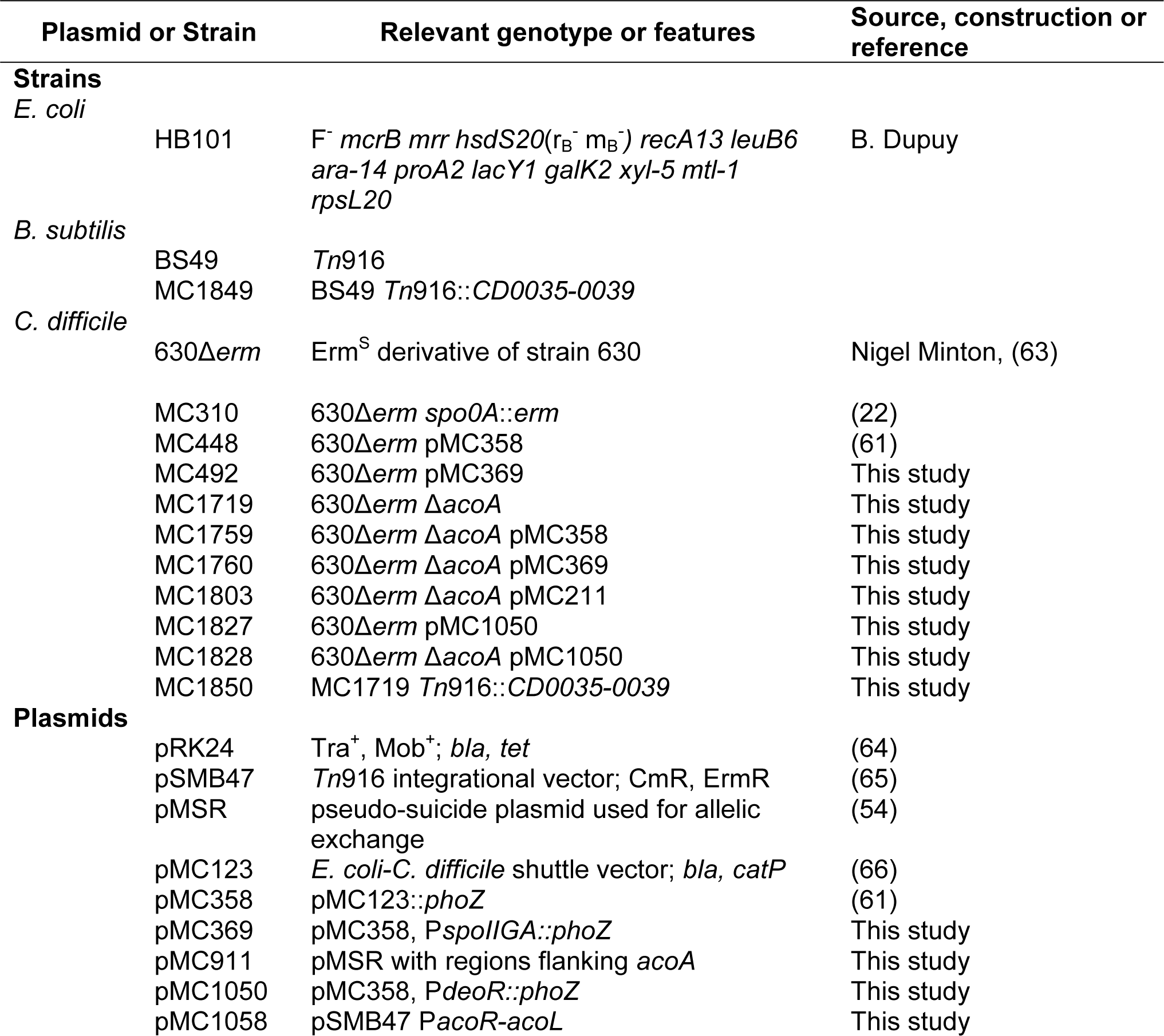
Bacterial Strains and plasmids.

| Plasmid or Strain |  | Relevant genotype or features | Source, construction or reference |
| --- | --- | --- | --- |
| **Strains** |  |  |  |
| *E. coli* |  |  |  |
|  | HB101 | F<sup>-</sup> *mcrB mrr hsdS20*(r<sub>B</sub><sup>-</sup> m<sub>B</sub><sup>-</sup>) *recA13 leuB6 ara-14 proA2 lacY1 galK2 xyl-5 mtl-1 rpsL20* | B. Dupuy |
| *B. subtilis* |  |  |  |
|  | BS49 | *Tn916* |  |
|  | MC1849 | BS49 *Tn916::CD0035-0039* |  |
| *C. difficile* |  |  |  |
|  | 630Δ*erm* | Erm<sup>S</sup> derivative of strain 630 | Nigel Minton, (63) |
|  | MC310 | 630Δ*erm spo0A::erm* | (22) |
|  | MC448 | 630Δ*erm* pMC358 | (61) |
|  | MC492 | 630Δ*erm* pMC369 | This study |
|  | MC1719 | 630Δ*erm* Δ*acoA* | This study |
|  | MC1759 | 630Δ*erm* Δ*acoA* pMC358 | This study |
|  | MC1760 | 630Δ*erm* Δ*acoA* pMC369 | This study |
|  | MC1803 | 630Δ*erm* Δ*acoA* pMC211 | This study |
|  | MC1827 | 630Δ*erm* pMC1050 | This study |
|  | MC1828 | 630Δ*erm* Δ*acoA* pMC1050 | This study |
|  | MC1850 | MC1719 *Tn916::CD0035-0039* | This study |
| **Plasmids** |  |  |  |
|  | pRK24 | Tra<sup>+</sup>, Mob<sup>+</sup>; *bla*, *tet* | (64) |
|  | pSMB47 | *Tn916* integrational vector; CmR, ErmR | (65) |
|  | pMSR | pseudo-suicide plasmid used for allelic exchange | (54) |
|  | pMC123 | *E. coli*-*C. difficile* shuttle vector; *bla*, *catP* | (66) |
|  | pMC358 | pMC123::*phoZ* | (61) |
|  | pMC369 | pMC358, *PspolIGA::phoZ* | This study |
|  | pMC911 | pMSR with regions flanking *acoA* | This study |
|  | pMC1050 | pMC358, *PdeoR::phoZ* | This study |
|  | pMC1058 | pSMB47 *PacoR-acoL* | This study |

### Strain and plasmid construction

The oligonucleotide primers used in this study are listed in **Table 2**. Primer design and the template for PCR reactions were based on *C. difficile* strain 630 (GenBank accession NC_009089.1). All plasmids were sequenced prior to use (GenScript). Details of plasmid construction are provided in the supplemental **Figure S7**. Plasmids were conjugated into *C. difficile* from *E. coli* or *B. subtilis* as previously described (52, 53). The *CD0036* (*acoA)* deletion mutant strain was created after introducing using the pseudo-suicide allelic exchange vector pMC911 and screening by PCR for loss of *acoA* (54). Complementation of the *acoA* mutant was achieved by integration of *Tn*196 carrying the full *acoRABCL* operon and promoter sequence from pMC1058, which was introduced into the *B. subtilis* strain BS49 by transformation (MC1849), and conjugated to MC1719, as previously described (53, 55).

**Table 2.**
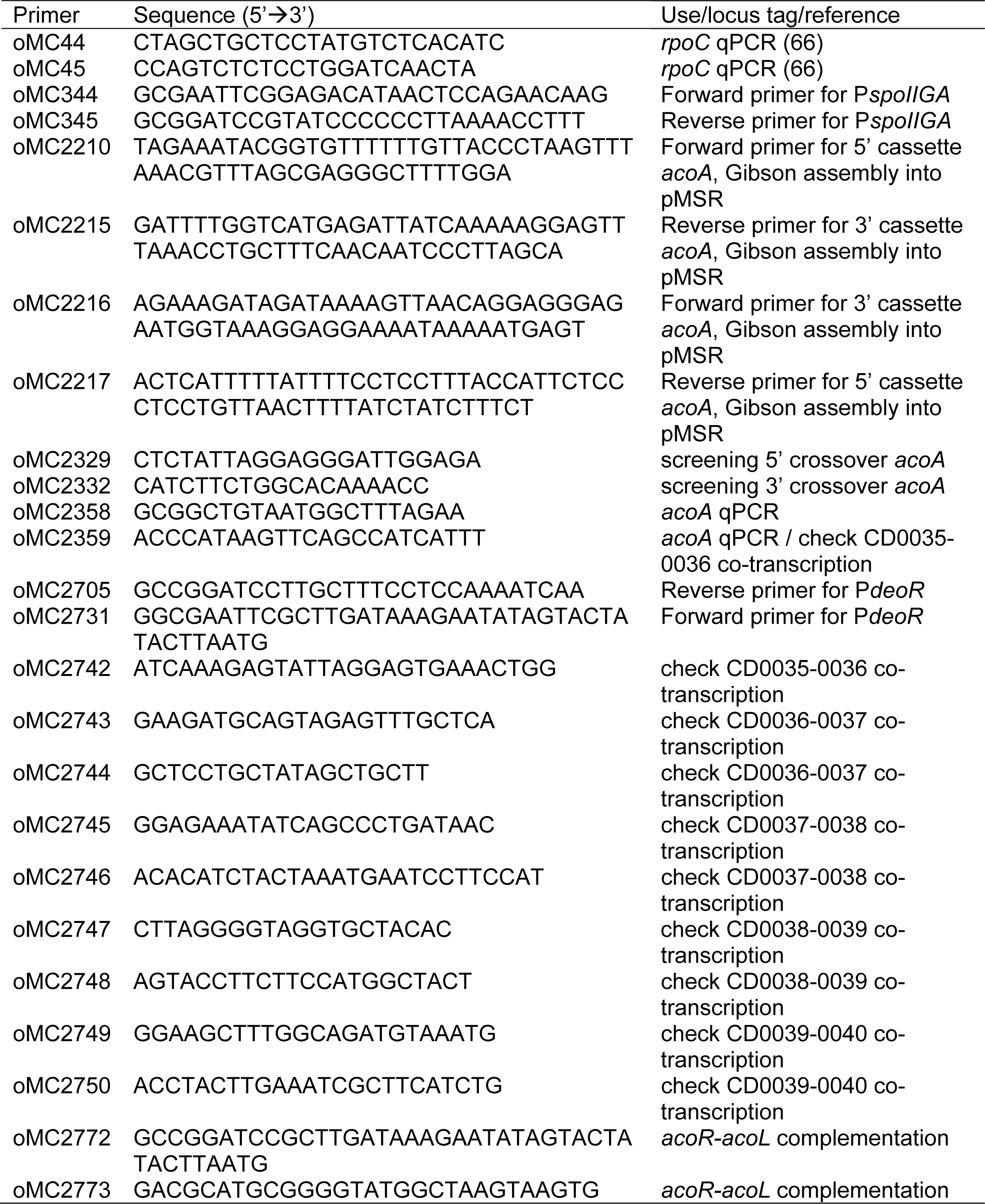
Oligonucleotides.

### Growth and sporulation in 70:30 medium

Sporulation assays were performed as previously described using a slightly modified 70:30 medium without the addition of Tris base and adjusted for pH, as shown (7, 56, 57). For sporulation in 70:30 broth, *C. difficile* was first cultured in BHIS with 0.1% taurocholate until mid-log phase, diluted in 70:30 to an OD_600_ of 0.3, and used to inoculate a 50 ml 70:30 culture (start OD_600_ of 0.03), adjusted to pH 7.2 or 6.2, respectively. Two hours after reaching an OD_600_ of 1 (time point T2), cultures were serially diluted and plated onto BHIS agar with 0.1% taurocholate for enumeration of total viable cells. After 24 h, culture samples were subjected to ethanol-resistance assays and spores enumerated as previously described (56).

For sporulation on 70:30 agar plates, log-phase BHIS cultures were diluted in BHIS to an OD_600_ of 0.5 and 250 µl of culture applied on 70:30 plates adjusted to diverse pHs, as indicated (56). After 24 h of growth, cells were scraped from plates, suspended in BHIS to an OD_600_ of 1, and evaluated for total cells and spores as previously described (7, 56). A *spo0A* mutant (MC310) was used as a negative control to ensure vegetative cell killing in ethanol resistance assays. The results represent three independent experiments and are presented as means with standard deviation of the means. Statistical significance was determined using a two-tailed Student’s *t*-test comparing the mutant to the parent strain or ANOVA, as indicated in the respective figure legends.

### Quantitative reverse transcription PCR analysis (qRT-PCR)

Cultures were grown on 70:30 agar and harvested from the plates twelve hours after inoculation (H12) into 6 ml ice-cold water:ethanol:acetone (3:1.5:1.5), and stored at - 70°C.

RNA isolation, DNase-I treatment (Ambion) and cDNA synthesis were performed as previously described (22, 26). Quantitative reverse-transcription PCR (qRT-PCR) with was performed with three technical replicates using 50 ng cDNA and the SensiFAST SYBR & Fluorescein Kit (Bioline) on a Roche Lightcycler 96 instrument. To confirm the absence of contaminating genomic DNA, cDNA synthesis reactions also included no reverse transcriptase for all samples. Results were calculated using the comparative cycle threshold method (58), normalizing expression to the internal control transcript, *rpoC*. For expression by pH condition, a one-way ANOVA and Dunnett’s test was performed for statistical comparison to the standard pH condition.

### Growth in minimal media

Growth curves were performed using a complete defined minimal media (CDMM) without the addition of D-glucose as previously described, (47) with slight modifications. Selenite (1 μM) (59) and zinc chloride (75 μM) (60), were added to the medium and the pH adjusted to 7.4 prior to filter sterilization. The CDMM base medium was supplemented with 30 mM acetoin (Sigma-Aldrich) or pyruvate (Fisher), as noted. Growth curves in minimal medium were carried out as follows: log-phase cultures were grown to an OD_600_ of 0.5 in BHIS medium, then diluted 10-fold into CDMM. Diluted cultures were used to inoculate CDMM medium broth at a 10-fold dilution for the growth assays to start at an OD_600_ of ∼0.01. Statistical significance was determined using a two-tailed Student’s *t*-test comparing the mutant to the parent strain.

### Alkaline phosphatase (AP) activity assays

*C. difficile* strains containing the transcriptional reporter fusions listed in **Table 1** were grown in 70:30 broth pH 7.2, with or without addition of 30 mM acetoin or pyruvate, as indicated. Cells were harvested at the indicated time points. AP assays were performed as previously described (61), without the use of chloroform for cell lysis. Technical duplicates for each strain and condition were averaged, and the results provided as the means and standard deviation for three biological replicates. A two-tailed Student’s *t*-test was used to compare the activity in the *acoA* mutant to the parent strain or a two-way ANOVA and Dunnett’s test was performed for statistical comparison of multiple strains and conditions, as appropriate.

### Detection of Toxins by ELISA

The *C. difficile* toxins TcdA and TcdB were quantified from the supernatants of cultures grown in TY broth (pH 7.4) for 24 h, according to the manufacturer instructions tgcBIOMICS (catalog no. TGC-E001-1). Technical duplicates were averaged and normalized per ml of cell culture. The results are presented as the means and standard deviations from three independent experiments. Statistical significance was determined using a two-tailed Student’s *t*-test comparing the mutant to the parent strain.

### Acetoin quantitation assay

A modified Voges-Proskauer test was used as a quantitative colorimetric assay to measure acetoin concentration in *C. difficile* supernatants from cultures grown in the absence or presence of 0.5 mM acetoin (38, 62). Briefly, active *C. difficile* cultures were diluted into duplicate BHIS cultures to an OD_600_ = 0.03. At OD_600_ = 0.5, 0.5 mM acetoin was added to one duplicate culture, and supernatants were harvested at this point (T0) and after six hours of growth (T6). A BHIS medium control with and without acetoin was included to control for acetoin stability. *C. difficile* supernatants were diluted 1:3 in distilled H_2_O. An acetoin standard curve was created in 0.25X BHIS. To a 96-well plate, 81 μl of the diluted culture supernatant or acetoin standard was added, followed by 56.7 μl of 0.5% creatine, 81 μl of 5% α-naphthol, and 81 μl of 40% KOH, mixing well after the addition of each reagent. After a 15 min incubation at RT, *A*_560_ was measured using a BioTek Synergy H1 plate reader.

## ACKNOWLEDGEMENTS

We give thanks to members of McBride lab for helpful suggestions and discussions during the course of this work. This research was supported by the U.S. National Institutes of Health through research grants AI116933 and AI156052 to S.M.M. The content of this manuscript is solely the responsibility of the authors and does not necessarily reflect the official views of the National Institutes of Health.

---

**Figure S1.**
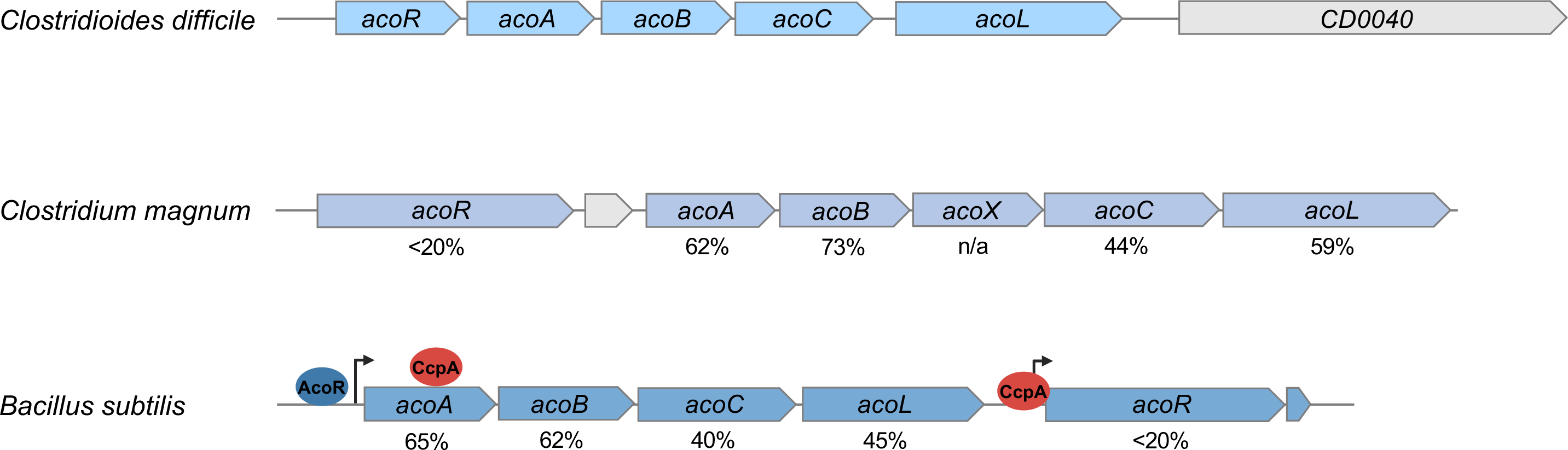
Comparison of acetoin gene clusters of *C. difficile* and other Gram-positive bacteria. Shown is the organization of the *aco* operons and accessory factors of *C. difficile* (AM180355), *C. magnum* (GCA_900129955.1), and *B. subtilis* (NC_000964). Numbers below the genes indicate the percent similarity of proteins to the corresponding *C. difficile* sequences. The approximate binding sites of the regulator CcpA (repressor), SigL, and AcoR (SigL- accessory activator)

**Figure S2.**
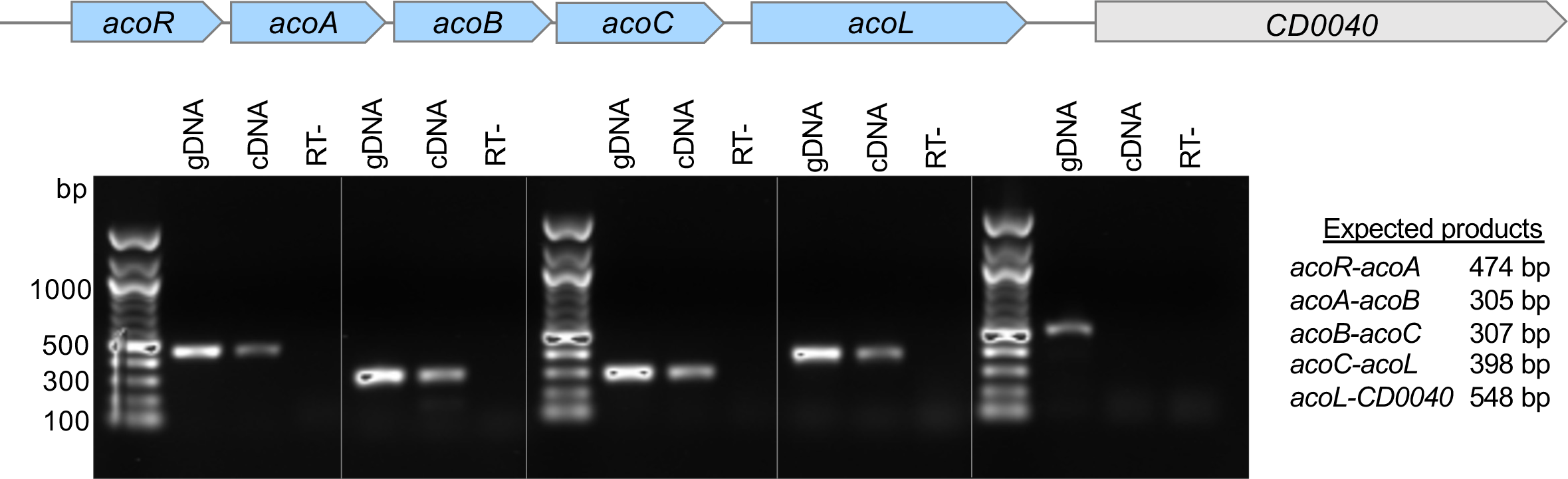
*CD0035 (acoR)* to *CD0039 (acoL)* are transcribed as an operon. Transcriptional units were examined by amplification of products from adjacent open reading frames. Cultures of 630&*erm* were grown in 70:30 medium to exponential growth phase (OD600 0.5) at pH 7.2, samples were collected for RNA and cDNA synthesis. PCR was performed using 50 ng genomic DNA (gDNA, positive control), 50 ng cDNA templates, or 50 ng cDNA without reverse transcriptase (RT-, negative control). Primer pairs include: *acoR-acoA,* oMC2742/oMC2359; *acoA-acoB,* oMC2743/oMC2744; *acoB-acoC,* oMC2745/oMC2746; *acoC-acoL,* oMC2747/oMC2748; and *acoL-CD0040,* oMC2749/oMC2750.

**Figure S3.**
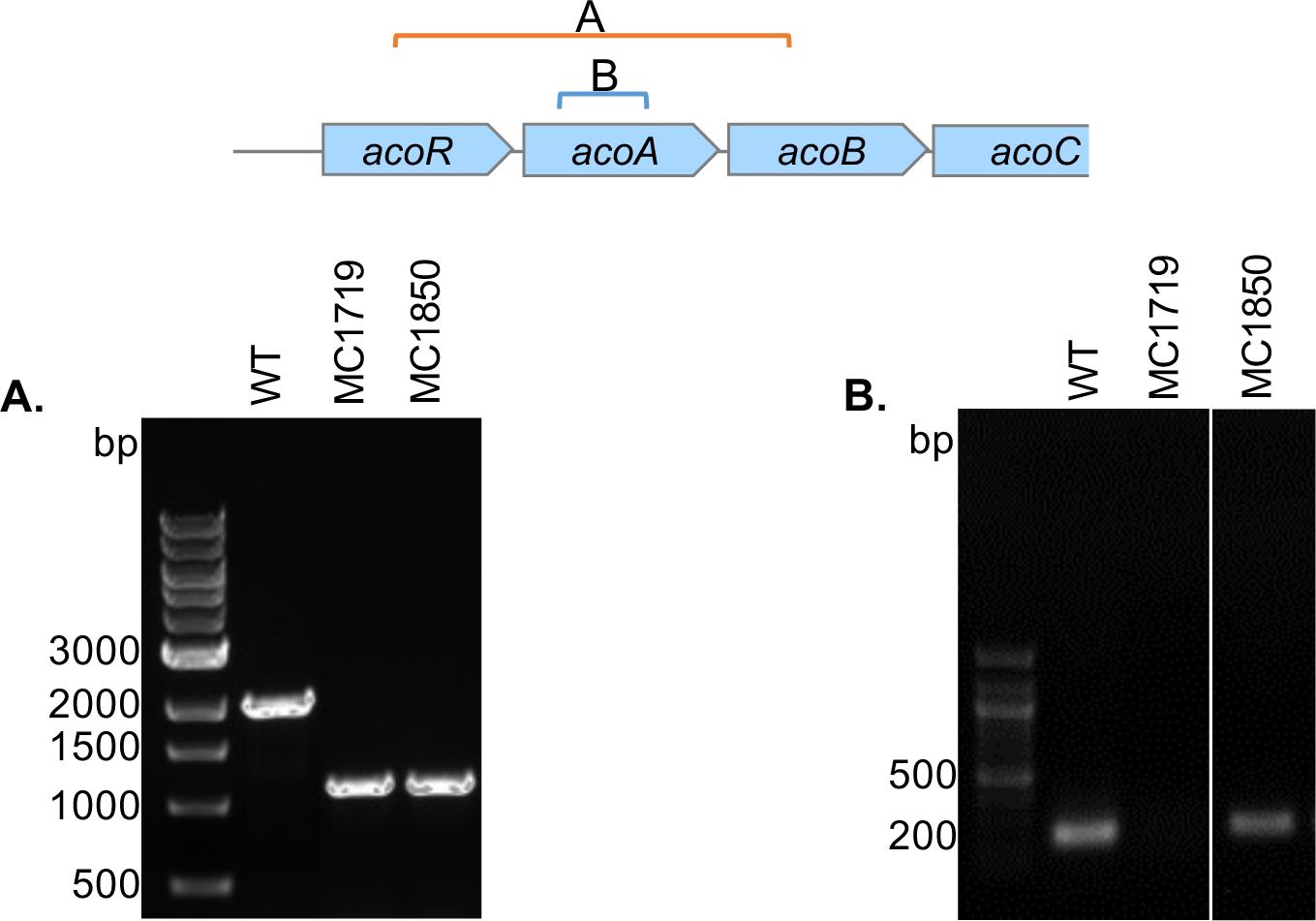
Confirmation of &*acoA* mutant and complemented strain. Genomic DNA from 630—*erm* (control, WT), &*acoA* (MC1719) or complemented &*acoA* Tn:*CD0035*-*0039* strain (MC1850) were used as templates PCR products were generated using primers **A)** flanking *acoA* (oMC2329/oMC2332; orange) to generate a wild-type product of 2125 bp and deletion product of 1156 bp. or **B)** primers within the *acoA* coding sequence (oMC2358/oMC2219; blue) to generate a wild-type product of 216 bp. Gels were cropped to show products only for relevant strains.

**Figure S4.**
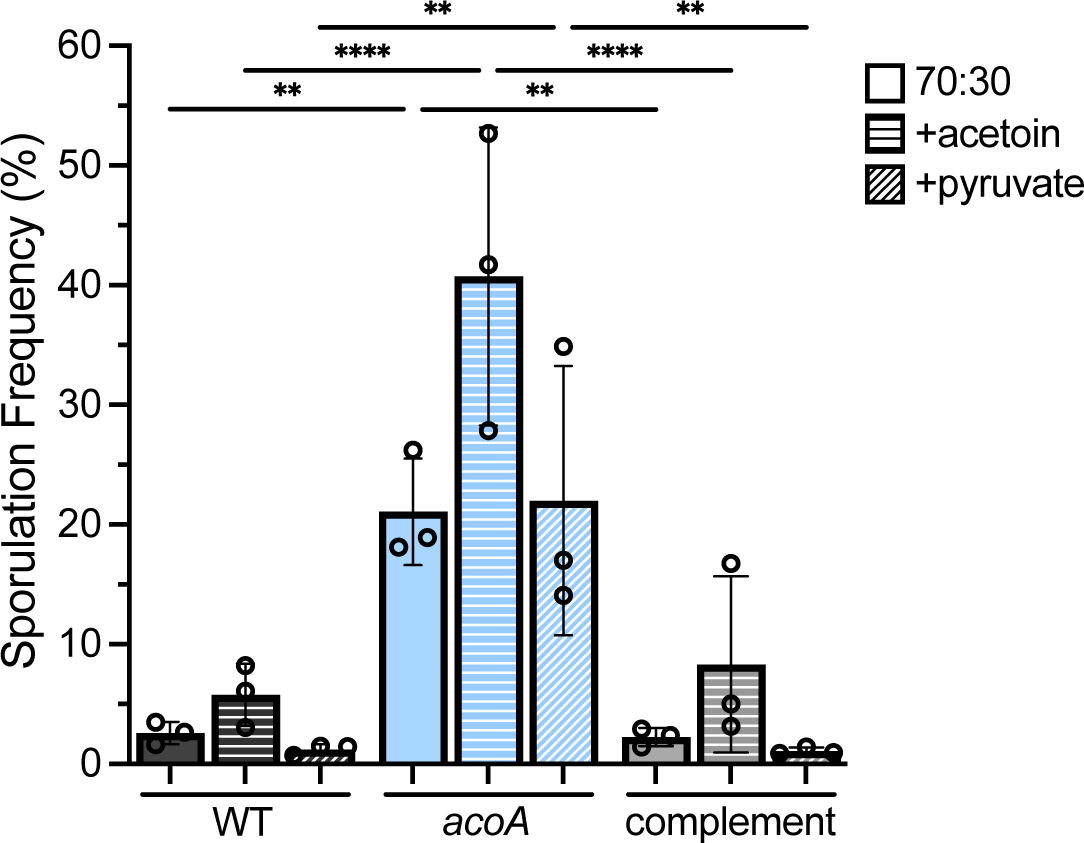
Complementation of sporulation in the *acoA* mutant. Ethanol-resistant spore formation of WT (630—*erm*), the *acoA* mutant (MC1719), and *acoA* complement strain (MC1850) grown in 70:30 broth culture (pH 7.2) for 24 h with or without 30 mM of acetoin or pyruvate, respectively. Sporulation frequency represents the mean percentage of spores relative to total viable cells at T2, shown with the standard deviation for three independent replicates. Data were analyzed by two-way ANOVA with Tukey9s post-hoc test. Asterisks indicate *P* values: * f 0.05; ** f 0.01; *** f 0.001; **** f 0.0001. No significant differences were identified between the WT and complemented strains.

**Figure S5.**
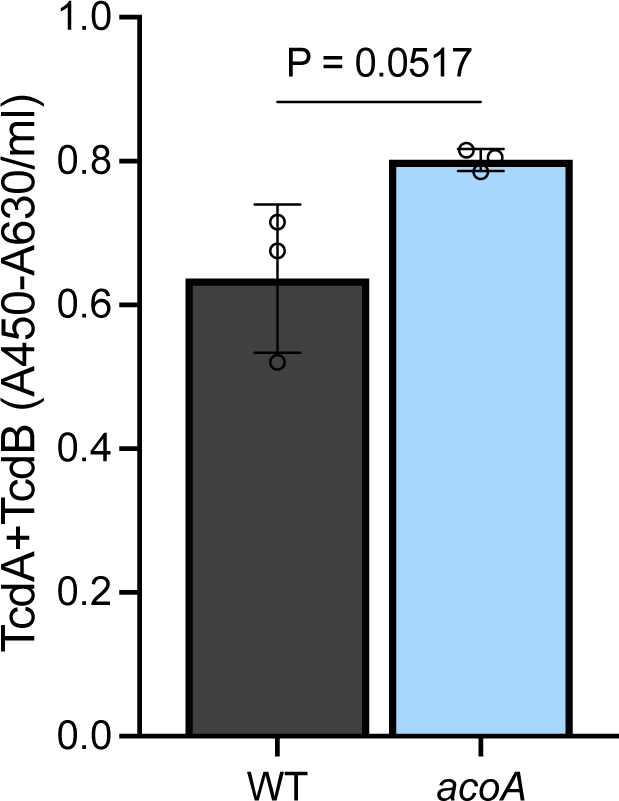
Toxin formation is not significantly impacted in the *acoA* mutant. Quantification of TcdA and TcdB present in the supernatants of WT (630—*erm*) and the *acoA* mutant (MC1719) grown in TY medium for 24 h and quantified by ELISA, as detailed in Materials and Methods. The means and standard deviation for three biological replicates are shown. Student9s unpaired *t*-test was used to compare the mutant to the parental strain.

**Figure S6.**
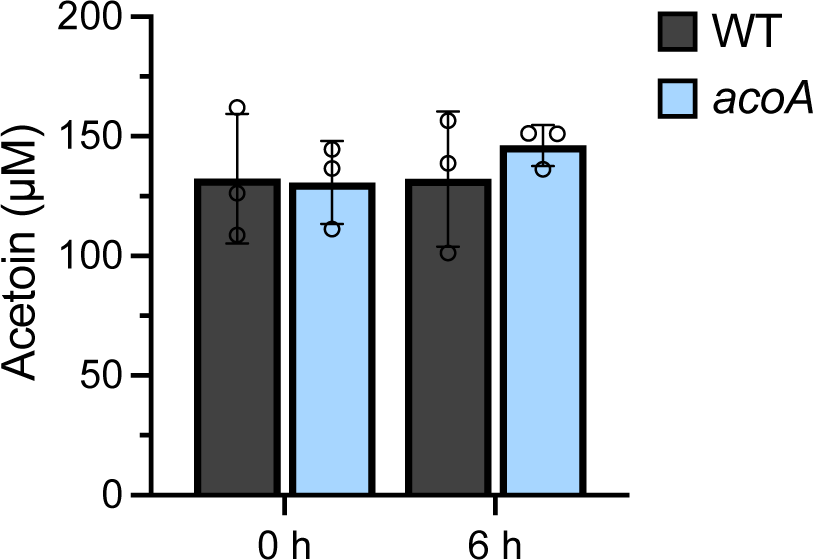
Acetoin utilization is comparable in WT and the *acoA* mutant. Acetoin concentration in supernatants from wild-type (630—*erm*) and the *acoA* mutant (MC1719) grown in BHIS broth with 0.5 mM acetoin. Supernatants were harvested from cultures during log phase following the addition of acetoin (0 h) and six hours later. No significant differences were observed.

**Figure S7. Plasmid construction** details *Plasmid construct details:*

**pMC369**: A 385 bp fragment containing the *spoIIG* promoter region was amplified using primers oMC344/oMC345 and cloned into pMC358 using the EcoRI and BamHI sites.

**pMC911**: A 507 bp fragment upstream of *acoA* was amplified using primers oMC2210/oMC2217 and a 511 bp fragment downstream of *acoA* was generated with primers oMC2215/2216. Both fragments were Gibson assembled together into pMSR at the PmeI site.

**pMC1050**: A 489 bp fragment upstream of *CD0035* (containing the *aco* promoter) was amplified using primers oMC2705/2731 and cloned using BamHI and EcoRI into pMC358.

**pMC1058**: The 6495 bp fragment containing the *acoRABCL* genes and upstream promoter region was cloned using BamHI and SphI into pSMB47.

